# Automatic classification of ICA components from infant EEG using MARA

**DOI:** 10.1101/2021.01.22.427809

**Authors:** I. Marriot Haresign, E. Phillips, M. Whitehorn, V. Noreika, E.J.H. Jones, V. Leong, S.V. Wass

## Abstract

Automated systems for identifying and removing non-neural ICA components are growing in popularity among adult EEG researchers. Infant EEG data differs in many ways from adult EEG data, but there exists almost no specific system for automated classification of source components from paediatric populations. Here, we adapt one of the most popular systems for adult ICA component classification for use with infant EEG data. Our adapted classifier significantly outperformed the original adult classifier on samples of naturalistic free play EEG data recorded from 10 to 12-month-old infants, achieving agreement rates with the manual classification of over 75% across two validation studies (n=44, n=25). Additionally, we examined both classifiers ability to remove stereotyped ocular artifact from a basic visual processing ERP dataset, compared to manual ICA data cleaning. Here the new classifier performed on level with expert manual cleaning and was again significantly better than the adult classifier at removing artifact whilst retaining a greater amount of genuine neural signal, operationalised through comparing ERP activations in time and space. Our new system (iMARA) offers developmental EEG researchers a flexible tool for automatic identification and removal of artifactual ICA components.

## 1. Introduction

The use of EEG in developmental cognitive neuroscience has led to a rich understanding of how the brain develops throughout early life. EEG has provided insights from birth into the development of skills such as face processing (e.g., Farroni, Csibra, Simion, and Johnson 2002) attention (e.g., Xie, Mallin and Richards, 2018), memory (e.g., Jones, Goodwin, Orekhova, Charman Dawson, Webb, and Johnson, 2020) and social interaction (e.g., Wass, Noreika, Georgieva, Clackson, Brightman, Nutbrown, Covarrubias, and Leong, 2018). It has also been pivotal in identifying risk factors associated with developmental disorders (e.g., Orekhova, Elsabbagh, Jones, Dawson, Charman, Johnson & the BASIS team, 2014) and later emerging psychopathology (e.g., Jones and Johnson, 2017). However, the field is challenged by a lack of scalable, standardised tools for artifact correction. In this paper, we present one lossless approach tuned for naturalistic artifact correction.

### 1.1. Traditional approaches to artifact removal

Despite its value, EEG recorded from paediatric populations is particularly susceptible to artifact contamination and typically contains fewer sections of clean uninterrupted data due to lower recording tolerances (Gabard-Durham, Leal, Wilkinson, and Levin, 2018; Debnah, Buzzel, Morales, Bowers, Leach, and Fox, 2020). One common approach to deal with this is to manually remove sections of the continuous data that are contaminated with artifact. However, this method of data cleaning can be problematic. For example, artifact correction for large EEG datasets can be very time consuming, and as developmental neuroscience is growing and EEG datasets are becoming larger, automated pre-processing tools are needed to efficiently process large-scale data, taking less time than manual cleaning (Webb, Bernier, Henderson, Johnson, Jones, Lerner, and Westerfield, 2015). Further manual cleaning is inherently subjective and there exist few comprehensive reviews to guide researchers (e.g., Chaumon, Bishop, and Busch, 2015). Recent studies have introduced methods for automatically identifying and removing segments of data contaminated by artifact in paediatric populations (e.g., Gabard-Durnham et al., 2018). These types of studies address the need for standardisation and speed but rely on complete removal of artifact-affected segments. Further, many of the currently available methods for paediatric EEG have procedures designed specifically for higher electrode density recordings, it is necessary to develop artifact correction approaches that are also flexible to low-density recordings, which are often used in infant EEG studies.

Recently, there has been a drive towards the use of more naturalistic paradigms in EEG research (Risko, Richardson, and Kingstone, 2016; Wass, Whitehorn, Marriott Haresign, Phillips, and Leong, 2020; Holleman, Hooge, Kemner, and Hessels, 2020). However, naturalistic EEG recordings provide additional analytical challenges over traditional screen-based tasks. For example, in traditional screen-based/ event-related tasks in which the child is passively exposed to a set of stimuli, artifacts are more randomly distributed with respect to stimulation. Removal of sections containing significant artifact can in this context be potentially beneficial, as visual experience during these sections might also be different (e.g., at its simplest the child might be fussing and not be attending to the image on the screen). However, in naturalistic paradigms, removal of whole sections of data is particularly problematic because data segments contaminated by artifact often covary with cognitive/ attentional processes of interest. Specifically, in naturalistic paradigms, the ‘stimulation’ is often child-controlled (e.g., the child turning to the parent in a naturalistic interaction), and so artifacts are more likely to be time-locked to neural signals of interest; the removal of artifact is thus likely to also affect the analysis of neural signals. Thus, we need approaches to the correction of artifact that remove artifactual signals from the EEG recording throughout the session, rather than removing whole segments of both signal and noise – so-called lossless pipelines.

### 1.2. Lossless approaches

Independent components analyses (ICA) applied to EEG data separates the contributing sources to the scalp EEG (Rutledge and Bouveresse 2013), which allows researchers to examine what mixture of pure source signals and their respective contributions make up each row of the data matrix (e.g data at each electrode) and to consider how these different source signals are weighted topographically (Makeig, Bell, Jung, and Sejnowski, 1996). By decomposing the EEG into its source components, researchers can inspect and remove components associated with artifact from the data and then remix the remaining components and project back into the original data format. This is a lossless approach as it does not involve the removal of entire sections of the data.

We note only one other attempt to provide a system for automatic ICA classification appropriate for paediatric EEG data. The adjusted-ADJUST program (Leach, Morales, Bowers, Buzzell, Debnath, Beall, and Fox, 2020) provides developmental researchers with an excellent framework for automating ICA classification from typical repeated stimulus EEG data. Leach and colleagues’ system achieved classification agreement with human coders of >85% with EEG recorded from 6-month-old infants. Whilst this is an impressive system, we feel that its application is limited for developmental EEG practitioners. Firstly, the adjusted-ADJUST program is set up to primarily deal with stereotypical eye movement artifact. Three of the five categories it sorts ICA-components into are related to ocular motor activity. Second, it is designed for event-locked paradigms with a repeated stimulus and is not able to incorporate continuous EEG data, such as the non-event locked paradigms, which are frequently used to study neural entrainment in parent-infant interactions (Wass et al., 2020). Third, validations of the system focused on the percentage of trials rejected, which further emphasises its suitability only for trial-based, pre-epoched data. In this manuscript we offer an alternative solution, that is flexible to continuous and event-locked data.

### 1.3. The MARA classification system

Many researchers perform manual classification to identify which of the components identified by ICA arise from genuine neural sources, and which are artifact. Other researchers have, however, attempted to automate this process. In this paper, we focus on one **automated** method, the Multiple Artifact Rejection Algorithm (MARA). This was originally designed for classification and rejection of non-neural/ artifactual ICA-components in adult EEG data (Winkler, Haufe, & Tangermann, 2011). The MARA classification system is grounded in the use of a binary linear classifier, whereby solving the equation (finding a separating

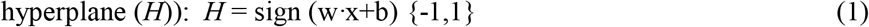

Where w is a weight vector obtained from samples of labelled training data, x is a feature vector and b is a bias term, classification of ICA- components as neural or artifact is achieved. The MARA classifier was originally trained using 690 ICA-components from an adult EEG reaction time study (n = 23 datasets), which had been manually classified as artifact/neural. The accuracy of the classifier was then tested on 1080 additional components from the same study. Accuracy was tested by comparing the results of the automatic ICA classification to manual ICA classification. The system achieved agreement rates of approximately 91%, (i.e., 9% of components were classified differently when comparing the automatic and manual classification). Accuracy was then further tested on new data from two other studies; an auditory event-related potential (ERP) paradigm (n=18 datasets); and a motor imagery BCI paradigm (n = 80 datasets), both with different channel setups and participants. Testing the performance of the classifier on the additional data revealed agreement/error rates between the automatic and manual classification of 85/15% (Winkler et al 2011).

Despite its popularity within adult EEG research, MARA has not received much attention within paediatric EEG research. This is perhaps because ICA itself is not widely used within traditional paediatric ERP research as a pre-processing tool. One previous study quantified the performance of MARA with paediatric EEG data. Gabard-Durnham and colleagues incorporated the classifier as part of their pre-processing tool kit (HAPPE- Gabard-Durnham et al., 2018), applying it to samples of high density (128 channels) resting-state EEG from infants and children aged 3-36 months. The authors reported extremely high rates of rejection (>85%) of ICA components when used as part of a conventional EEG pre-processing pipeline (e.g., including referencing, filtering, channel rejection/interpolation and trial/ continuous data rejection). The lack of objective tools for automated EEG preprocessing is a problem as developmental cognitive neuroscience is growing in scale and robustness (Debnah et al., 2020; Desjardins, Van Noordt, Huberty, Segalowitz, and Elsabbagh, 2020). In the present study, we aim to address this need for systems for automatic ICA cleaning of infant EEG data that can be incorporated among other standard pre-processing procedures.

### 1.4. The need to tune artifact-removal approaches to infant EEG data

Infant EEG has unique properties, requiring the design of specific tools for processing. EEG recorded from infants differs from that of children (Lepage, Jean◻François, and Théoret, 2006) and adults (Strogenova, Orekhova, and Posikera, 1999). For example, the canonical frequency bands e.g., delta (1-4Hz), theta (4-8Hz), alpha (9-13Hz) etc observed in adult EEG are observed at lower frequencies in infant EEG (Orekhova, Stroganova, Posikera, and Elam, 2006). Peaks in the power density spectrum that are associated with alpha activity typically observed in the 9-13Hz range in adults can be seen clearly between 6 and 9Hz in one-year-old infants (Strogenova et al., 1999) and are lower still in younger infants (Marshall, Bar’Haim, and Fox 2002). We also know that infant EEG tends to show greater power at lower (<6Hz) frequencies and that during development there is an observable increase in power at higher frequencies (Marshall et al., 2002). Whilst these differences have been observed in scalp level EEG data and not at a source level, this evidence highlights differences in the distribution of power at lower frequencies and the overall composition of the 1/f power density curve for infant vs adult EEG.

There is also evidence to suggest that the spatial properties of infant EEG differ from those typical of adult EEG. For example, we know that infant alpha activity projected onto central scalp electrodes is present only in later stages of infant development, presumably accompanying advances in motor skills (Cuevas, Cannon, Yoo, and Fox, 2014), although the sources of these scalp activations are yet to be identified. Further, at the source level infant EEG is often more bilaterally symmetrical than adults (Piazza, Cantiani, Miyakoshi, Riva, Molteni, Reni, and Makeig, 2020), although strong spatial asymmetry or localisation to a specific spatial point can be a good indication of artifactual source components (Chaumon, et al., 2015). This evidence highlights that infant EEG source components do contain spatially distinct properties to those of typical adult EEG. Overall, the evidence highlights the differences between adult and infant EEG data both at the scalp and source level. It should be clear from reviewing these studies that attempting to classify infant ICA components using training data from adult EEG would lead to sub-optimal results.

### 1.5. Current study: motivation and goals

In this study, we examine the performance of MARA when applied to samples of 32-channel infant EEG data acquired during naturalistic social interactions. We then adapt the MARA system to better fit the characteristics of infant EEG data. We do this in two ways; (1) by adapting the relevant time-frequency properties derived from the ICA used in classification; (2) by retraining the base classifier using data from infant EEG recordings. From here on we refer to the retrained classifier as iMARA.

To validate the performance of iMARA, we first looked at the inter-rater agreement of ICA components between three expert hand coders. Specifically, we looked at inter agreement between expert manual coders by calculating the Mean Square Error (MSE) on an n=15 subsample of infant and adult ICA-components from dataset 1. We then compared MARA and iMARA to the validated manually labelled infant ICA components across two validation studies: first (classifier validation 1), we tested the two classifiers’ agreement with ICA-components manually classified by rater 1 on the full n=44, 1180 component dataset (again using MSE). Second (classifier validation 2), we tested the two classifiers’ performance on infant ICA components from an unseen dataset (n=25, 670 components) obtained during a different recording session. In our final validation (classifier validation 3), we looked at ERP data generated using different methods to examine in greater detail their ability to remove specific types of artifact. In validation 3, differences in performance were quantified by comparing peak amplitude potentials across four conditions: i) data cleaned using iMARA; ii) data cleaned using MARA; iii) data cleaned using manual classification; iv) data not cleaned using ICA (‘raw’).

Evidence from co-registered EEG and eye-tracking studies using free viewing experimental paradigms has shown that when visual responses (e.g. a stimulus appearing on-screen) co-vary with eye movements (e.g. horizontal/ vertical saccades) separation of these signals is possible based on their time and spatial properties (Plöchl, Ossandon and Konig, 2012). For example, some types of eye movement artifacts e.g., vertical and horizontal eye movement transients (i.e., only lasting ~200ms) peak at ~100ms post saccade onset and have anteriorly dominated topographies, whereas visual components tend to peak 100-200ms after the peak of the artifact and have occipitally dominated topographies (Plöchl et al., 2012). Based on these findings and inspection of our data time-locked to saccade onsets, we set up our comparison between the four cleaning methods described above as follows. For comparison of removal of eye movement artifact time-locked to saccade onset, we compared peak amplitudes of potentials over frontal pole electrodes (FP1, FP2, AF3 and AF4) in the time window −100 (saccade onset) to 100ms (see also figure 3 for visual representation). For comparison of retention of visual response (i.e., the neural signal of interest) we compared peak amplitudes of potentials in the 200-300ms time window over occipital electrodes (PO3, PO4, O1, O2, O3). We also compared amplitudes in the 200-300ms time window over central electrodes (C3, CP1, CP5, CP6, CP2, C4, Cz) to examine how these signals propagated across the scalp.

## 2. Methods

### 2.1. Ethics statement

This study was approved by the Psychology Research Ethics Committee at the University of East London. Participants were given a £50 shopping voucher for taking part in the project.

### 2.2. Participants

The same experimental paradigm was used for all validation dataset’s, but recordings were taken from different sessions (weekly sessions 1 and 8 as part of a broader, 8-week programme of research).

Dataset 1 (Validation 1), 44 healthy (23 F, 21 M) infants participated in the study along with their mothers. Infants were aged mean 10.72 months, std=1.31. Dataset 1 was taken from the infant’s visit 1 data.

Dataset 2 (Validation 2), 25 healthy (12 F, 13 M) infants contributed data. Infants were aged mean 12.60 months, std=1.27. Dataset 2 included the same infants with data taken from visit 8.

Dataset 3 (Validation 3), 36 healthy (17 F, 18 M) infants contributed data. Infants were aged 10-12 months (mean 10.70 months, std = 1.08). Dataset 3 is a subset of dataset 1.

### 2.3. Experimental set-up and procedure

Infants were positioned immediately in front of a table in a highchair. Adults were positioned on the opposite side of the 65cm-wide table, facing the infant. Adults were given toys to play with across a tabletop and asked to “play with their infant as they would normally do at home”. Adults were also asked to lower the volume of their vocalisations to reduce the level of speech-related contamination in the EEG. Dual EEG was continuously acquired from the parents and infants for the approx. 25 min duration of the play session. For this study, we used only the infant’s EEG.

### 2.4. EEG data acquisition

EEG signals were obtained using a dual 32-channel Biosemi system (10-20 standard layout). EEG was recorded at 512 Hz with no online filtering using the Actiview software.

### 2.5. EEG artifact rejection and pre-processing

A fully automatic artifact rejection procedure was adopted, following procedures from commonly used toolboxes for EEG pre-processing in adults (Mullen 2012; Bigdely-Shamlo, et al., 2015) and infants (Gabard-Durham et al., 2018; Debnath et al., 2020). This was composed of the following steps: first, EEG data were high-pass filtered at 1Hz (FIR filter with a Hamming window applied: order 3381 and 0.25/ 25% transition slope, passband edge of 1hz and a cutoff frequency at −6db of 0.75hz). Although there is debate over the appropriateness of high pass filters when measuring ERP’s (see Widmann and Schröger, 2012), we aimed to obtain the best possible ICA decomposition. The parameters we used were set up following recent work (e.g., Dimigen 2020) that examined the removal of eye movement artifacts from free viewing EEG using ICA. Second, line noise was eliminated using the EEGLAB (Delorme and Makeig 2004) function clean_line.m (Mullen 2012). Third, the data were referenced to a robust average reference (as described in Bigdely-Shamlo et al., 2015). Fourth, noisy channels were rejected, using the EEGLAB function pop_rejchan.m. Fifth, the channels identified in the previous stage were then interpolated back, using the EEGLAB function eeg_interp.m. The mean number of channels interpolated in this way was 4.2. In some datasets, channel interpolation reduced the overall rank of the data leading to a fewer number of components than channels as is the norm with ICA. Interpolation is commonly carried out either before or after ICA cleaning but in general, has been shown to make little difference to the overall decomposition (Delorme and Makeig 2004). Sixth, the data were low-pass filtered at 20Hz, again using an FIR filter with a Hamming window applied identically to the high-pass filter. (In the SM we also report a comparative analysis in which data were low pass filtered at 40Hz instead of 20Hz (see SM section 1.5)). Seventh, continuous data were automatically rejected in a sliding 1s epoch based on the percentage of channels (set here at 70% of channels) that exceed 5 standard deviations of the mean channel EEG power. For example, if more than 70% of channels in a given 1-sec epoch exceed 5 times the standard deviation of the mean power for all channels then this epoch is marked for rejection. This step was applied very coarsely to remove only the very worst sections of data (where almost all channels were affected), which can arise during times when infants fuss or pull the caps. This step was applied at this point in the pipeline so that these sections of data were not inputted into the ICA. The average amount of data retained in this way was 88% (std 0.1). Data were then concatenated and ICAs were computed on the continuous data using the EEGLAB function runica.m. The mean amount of data entered into the ICA was 21.2 minutes. In the raw data condition, we followed the same procedure but without any ICA correction.

### 2.6. Video coding

Video recordings were made using Canon LEGRIA HF R806 camcorders recording at 50fps positioned next to the child and parent respectively. Video recordings of the play sessions were coded offline, frame by frame, at 50 fps. This equates one frame to a maximum temporal accuracy of ~20ms. Coding of the infant’s gaze was performed by two independent coders. Cohen’s kappa between coders was >85%, which is high (McHugh, 2012). For our ERP analysis, EEG was time-locked to the onset of gaze/ saccade offline based on the video coding using synchronized LED and TTL pulses.

### 2.7. Hand identification of components for the training set

A full description of how components were identified as either neural or artifactual by human coders is given in Appendix A. Briefly, components were judged first on their topography, second on their power spectrum, and third on their time course, using similar principles to those suggested for adult EEG data (e.g., Chaumon et al, 2015). Components were marked as artifact/ rejected only under the null hypothesis – which in this case is that the component is not considered to contain notable amounts of the neural signal. Where a researcher was in doubt over whether a component contains real EEG (neural) we opted to retain that component.

### 2.8. Inter expert reliability

As within any classification system, performance is measured concerning a criterion representing ‘true value’ or ‘perfect classification’. There exists no gold standard upon which to test any classifiers performance. As manual classification is the typical approach for ICA data correction (Chaumon et al., 2015) and has been used as a platform to test automatic classification in previous studies (Winkler et al., 2011), we tested the MARA and iMARA systems performance against manual ICA classification. To validate our manual coding we asked 3 experts to independently rate ICA-components from infant and adult EEG data (see Table S1). We examined whether similar levels of agreement between coders could be achieved for infants ICA components as compared to those in adult data. Results are reported in section 3.1. Previous research using automated classification methods with adult data from screen-based tasks have reported an error in inter expert agreement levels of ~10-13% MSE (Winkler et al., 2011).

The measure of performance we use in this study is mean square error (MSE), as has been used in previous automatic classification studies (Halder, Bensch, Mellinger, Bogdan, Kübler, Birbaumer, and Rosenstiel, 2007; Winkler et al., 2011). In its simplest interpretation, MSE is a measure of error rate between systems. For example, an MSE of 0.25 would indicate that the automatic and manual classifiers differed on 25% of the components examined.

### 2.9. Set-up and paradigm for validation dataset 3 (ERP Analysis)

To further test the performance of the different classifiers, we contrasted the different systems’ ability to remove stereotypical artifact from an ERP analysis. This analysis examines event-locked changes relative to infants’ spontaneous gaze shifts during a free-flowing naturalistic interaction. Specifically, we examined moments where infants shifted from looking at a puppet, held at the same height as their mother’s face, c.10° from the midline (counterbalanced between left and right) to looking at their mother’s face, who was always positioned directly in front of the infant. To boost trial count, we concatenated epochs of gaze shifts when the adult was already looking at their infant (i.e., the infant looks to direct gaze) and when the adult was looking at the puppet (i.e., the infant looks to averted gaze) as both are time-locked to a shift in infant attention. For this analysis, we extracted epochs from the continuous data that are time-locked (time 0) to the infant’s fixation onset (saccade onset at −100ms). Epochs were taken from 1.5s before the fixation onset to 1.5s after. Mean (39.4) std (12.9) gaze shifts were included per participant.

ERPs were compared over frontal, central and occipital scalp regions. Details of which electrodes were used in each cluster can be found in the supplementary materials section (SM section 1.3, Table S3). We compared the peak amplitudes of potentials over frontal pole electrodes in the time window −100 (saccade onset) to 100ms (following Plöchl et al., 2012). For comparison of retention of visual response (i.e., the neural signal of interest), we compared peak amplitudes of potentials in the 200-300ms time window over occipital electrodes. We also compared amplitudes in the 200-300ms time window over central electrodes to examine how these signals propagated across the scalp. These comparisons were repeated for all four methods of data cleaning (e.g., iMARA, MARA, manual cleaning, and raw).

Differences in peak amplitude were quantified using the adaptive mean approach. This process involves identifying the peak latency of the ERP potential on a subject-by-subject basis using a broad (100ms) time window, centred around the time window of interest. For example, in our analysis, we were interested in activity in the −100 to 100ms time window. In this case, the adaptive mean approach looks for the latency of the data point with the maximum amplitude +/− 50ms around the centre of the time window (0ms). Once the peak latency has been identified we took an average of the activity in a 20ms window around the peak (e.g. as described by Hoorman, Falkenstein, Schwarzenau, and Hohnsbein 1998). This approach is preferred over the more basic comparison of absolute peak amplitudes which would be more susceptible to spurious noise spikes and/or unrepresentative data (Cohen, 2014). All ERP data were baseline corrected using data from the time window −1000 to −700ms pre gaze onset.

### 2.10. The MARA system for automatic classification of neural/ artifactual components

The MARA classification system identifies artifactual source components from samples of EEG data. For a detailed explanation and the original source code, please refer to (https://irenne.github.io/artifacts/). In brief, Winkler and colleagues (2011) trained a binary linear classifier to separate neural and artifactual ICA decompositions based on a training dataset of manually labelled ICA components. The comparison between neural and artifactual components was conducted by examining six features derived from the ICA time-frequency properties (see Figure 1).

**Fig. 1.**
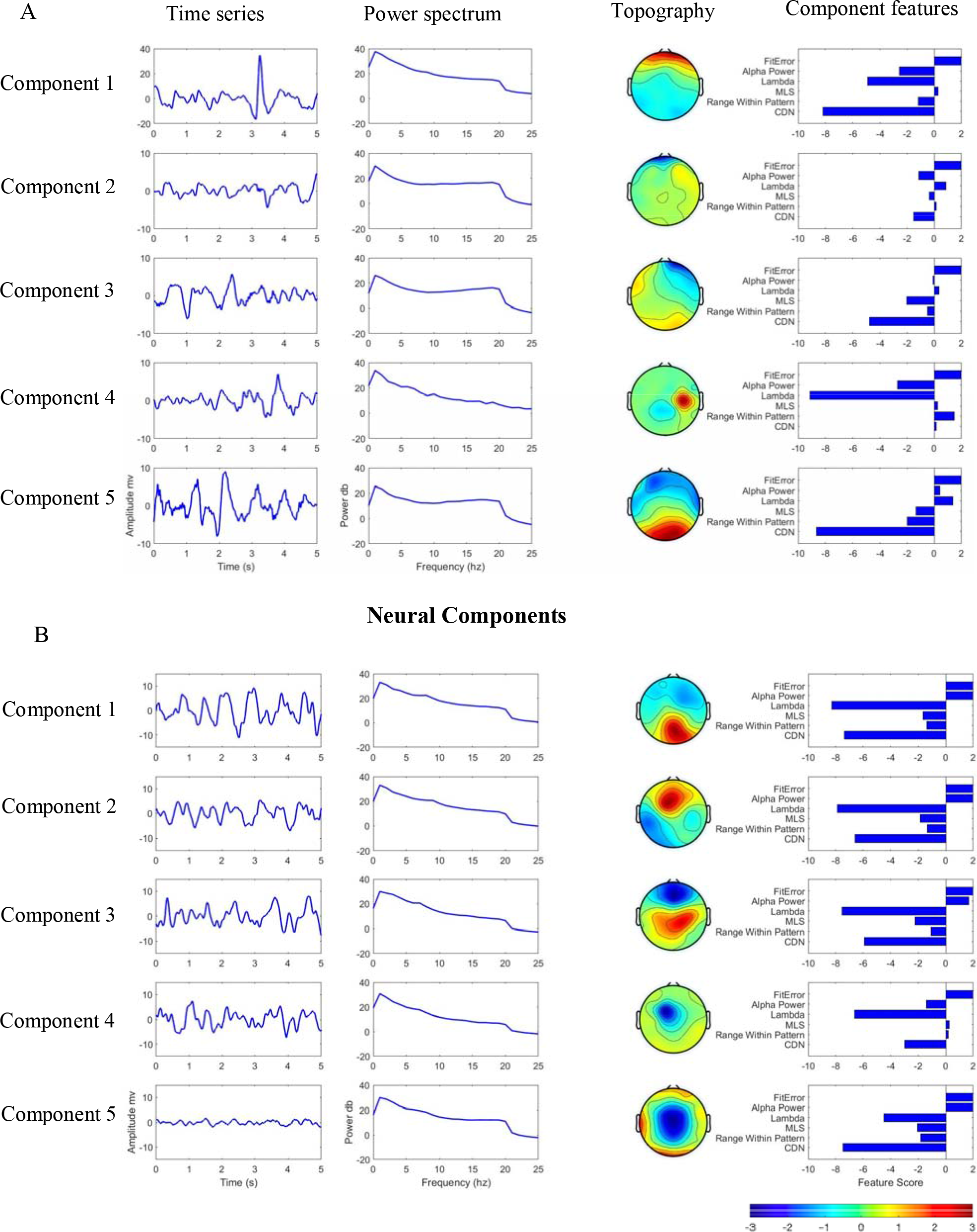
Examples (taken from the present study) of artifactual and neural ICA components identified by iMARA. A) Examples of components identified as artifact by iMARA. B) Examples of components identified as neural by iMARA. For both, the first column shows five-second segments of the component’s time course; the second shows the component power spectral density; the third shows the topographical activations; and the fourth their scores for the six features used in classification. Detailed descriptions of the six features are given in section 2.11.

**Fig. 2.**
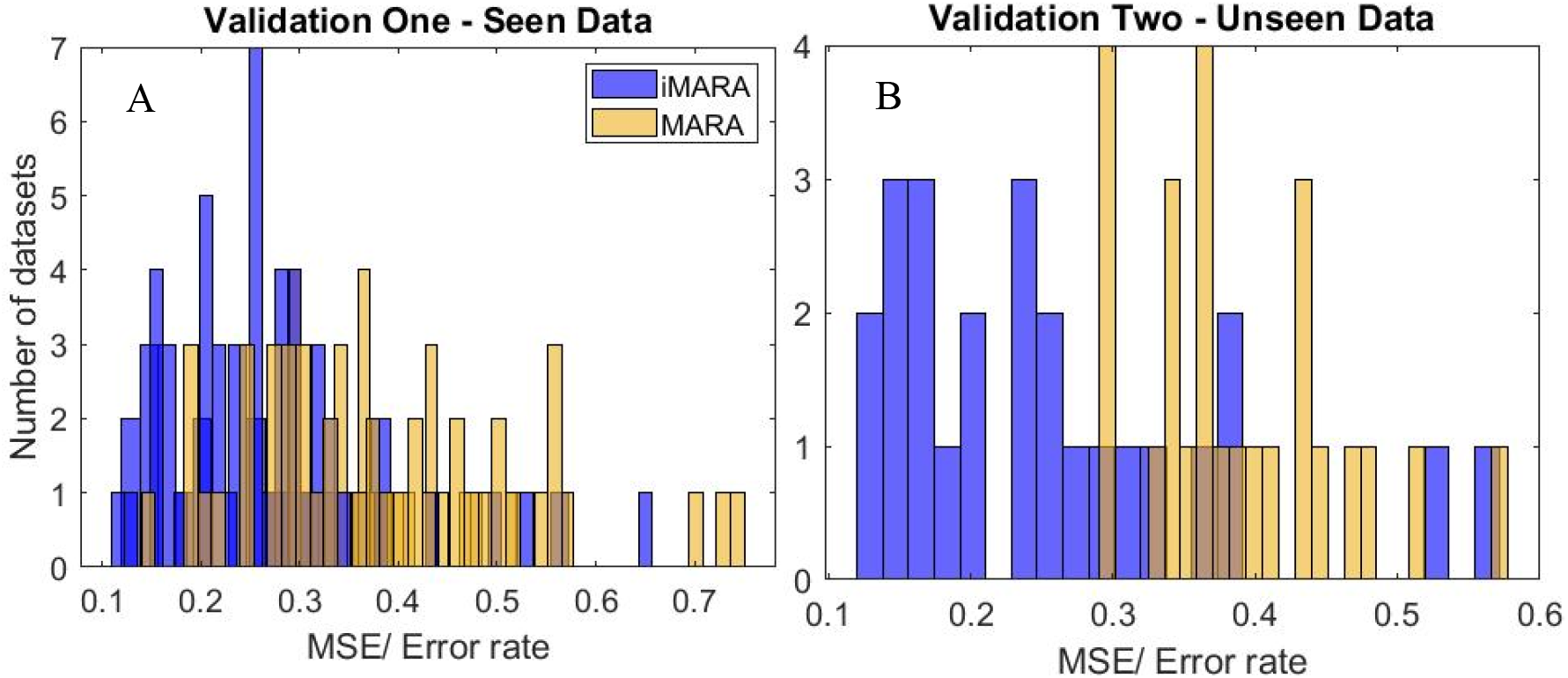
Classification performance for original (MARA) and retrained (iMARA) systems on ‘seen’ and ‘unseen’ data. A) Mean Squared Error (MSE) between original (‘MARA’ - yellow) and retrained (‘iMARA’ - blue) classifiers and manually classified ICA components for validation one (seen data) for each participant (n=44) of dataset one. B) MSE between iMARA/MARA and the manual classification for validation two (blind data) for each participant (n=25) of dataset two.

### 2.11. Feature selection

In the original paper, the following six features were selected for use in the MARA system. These were originally chosen through an embedded feature selection process (e.g., integrated as part of the learning algorithm) whereby the authors obtained rankings of importance/ effectiveness of 38 different time/frequency/spatial features of the data (for more details see Winkler et al., 2011). This revealed that inclusion of additional features (beyond the six included) did not increase classification performance.

The following two features relate to the component spatial distribution:

<u>Current Density Norm (CDN)</u> – estimation of source position of a component concerning x,y,z spatial coordinates. This process involves dipole fitting the source components (using functionality contained with EEGLAB) and applying an appropriate forward head model (we considered 2142 locations arranged in a 1 cm spaced 3D-grid) and seeking the source distribution with minimal l2-norm (i.e., the ‘simplest’ solution, Winkler et al., 2014).

Components with a high CDN indicate likely artifact. For example, it can be seen that on Figure 1a) component two, three and four all have a relatively high CDN score. These can be compared with components one, two and three of Figure 1b, which all have a relatively low CDN score and were classified as neural. This feature was unchanged from the original study.

<u>Range Within Pattern</u> – the absolute difference between the minimum and maximum of a component’s pattern (spatial distribution) - i.e., how localized the activation is to one position/ electrode. Comparing components two and four in Figure 1a and 1b, we see that artifactual components have a relatively higher range within pattern indicating that these sources are more localized to a singular point, which is taken as an indication of an artifactual component to the classifier. This feature can arise, for example, from poor contact between the surface of the electrode and the scalp. This feature was unchanged from the original study.

The following two features relate to the component-time series:

<u>Mean Local Skewness (MLS)</u> – the mean absolute local skewness of an ICA-component time series, taken in a 1 and 15s (two separate features) sliding time window and then averaged. The idea being that blink components for example would contain epochs with very high amplitude data. This data would be more skewed than a typical alpha generator in which you would expect amplitude to be comparatively unchanged across epochs.

For example comparing components one, two and three in Figure 1a and 1b, we see that a relatively high MLS indicates artifact, as this component’s time series might contain more high amplitude noise spikes than components with a low MLS. High MLS might arise from faulty electrodes, but is also an indication of an ocular motor artifact. For example, in Figure 1a component one, a stereotypical blink component has a relatively high MLS and contains frequent high amplitude spikes in the time series. This feature was unchanged from the original study.

The following two features relate to the component spectral distribution:

<u>Lambda and Fit Error</u>- the deviation of a components power spectrum from a pseudo 1/frequency curve, created by three points of the log spectrum: (1) value at 2 Hz, (2) local minimum in the band 5-13 Hz, (3) local minimum in the band 33-39 Hz. The spectrum of muscle artifacts, characterized by unusually high values in the 20-50 Hz range, is thus approximated by a comparatively steep curve with high lambda and low fit error. Lambda and fit error are independent features; whereas lambda is a measure of the deviation from the pseudo curve just in the alpha and beta ranges (i.e., steepness of transition between the two), fit error is a measure of the deviation of the components 1/f curve from the entire pseudo 1/f curve between delta to beta.

For example from component two in Figure 1a and 1b, we can see that low lambda (i.e., a less steep curve between alpha and beta) indicates a neural component, whereas high lambda (i.e., a steeper upward curve between alpha and beta) indicates artifact. We can also see that fit error does not always distinguish well between neural and artifactual components in these examples. This is because a neural component with a high alpha peak and an artifact component with a steep upward curve between alpha and beta would both give a high fit error, which can make classification using fit error alone difficult. We adjusted the frequency features to better fit the characteristics of infant EEG data. For fit error instead of taking values at 2hz, 5-13hz and 33-39hz as used in MARA, we take values at 2hz, 5-9hz and 12-19hz. Further for lamda instead of comparing activity in the 8-15hz range to the pseudo 1/f curve as used in MARA, we compared activity in the 6-13Hz to the pseudo 1/f curve.

<u>Alpha Power</u> – The average log band power of the alpha band (8–13 Hz).

From components one and four in Figure 1a and 1b, we can see that high alpha band power indicates a neural component, whereas low alpha power indicates artifact. Instead of taking a value for alpha power in the 8-13hz range as used in MARA, we take a value for alpha power in the 6-9hz range.

We retrained the MARA system using 617 ICA components from infant EEG data taken from dataset 1 (n=25 datasets, each contributing on average 25 ICA components) and using the six features, with the amendments that we have described above.

## 3. Results

First (section 3.1) we validated our manual classification by comparing it with manual classification from two other independent experts. Then, we perform three validation studies to test the performance of iMARA on infant data: first (classifier validation 1, section 3.2), we tested iMARA and MARA’s agreement with manually classified ICA-components by rater 1. Second (classifier validation 2, section 3.3), we test iMARA and MARA’s performance on ICA components from an unseen dataset. Third (classifier validation 3, section 3.4), we examine ERP data generated using the different methods to examine in greater detail their ability to remove specific types of artifact.

### 3.1. Inter-rater validation

To first validate our coding, we asked three experts independently to classify random subsamples of infant (n=15 datasets, average 25.6 ICs, taken from dataset 1) and adult (n=15 datasets, average 28.4 ICs, taken from dataset 1) EEG data. Full comparison details are given in SM section 1.2, Table S2. Between the 3 experts, the average disagreement rate for infant data was 18% (range across three all three experts 14-22%), whereas for adult data it was 15% (range across three experts 12-18%), which is in line with previous reports of human-human error rates for adult EEG data of 10-13% (e.g., Winkler et al., 2011). An independent sample test revealed no significant differences in the average agreement between adult and infant ICA-components t(14) = 0.98, p = 0.42.

### 3.2. Classifier validation 1

We tested the retrained classifiers performance against manually classified ICA components from validation dataset 1. This resulted in an averaged MSE between iMARA and the manual classification of 26.59% (sd = 9.93%, range = 54.11%). In comparison, when using the original MARA training data and the original feature extraction routine on dataset 1, the MARA classifier performed with an MSE of 38.35% (sd=15.01%, range = 60.19%). A paired samples t-test comparing the percentage of correctly identified components from validation dataset 1 for iMARA vs MARA indicated that MARA had a significantly lower level of agreement with the manual classification than iMARA t(43) = −5.94, p = <0.01. The effect size for this analysis was *d*=0.92.

### 3.4. Classifier validation 2

We then tested iMARA on an unseen dataset (dataset 2). Classification of the (645) unseen components led to an averaged MSE between iMARA and manual classification of 24.80% (std=8.22%, range=55.43%). In comparison, MARA performed with an MSE of 38.13% (std=8.12%, range=26.63%). A paired samples t-test comparing the percentage of correctly identified components from validation dataset 2 for iMARA vs MARA indicated that the original MARA had a significantly lower level of agreement with manually classified ICA components than iMARA t(24) = −4.50, p = <0.01. The effect size for this analysis was *d*=1.63.

### 3.4. Classifier validation 3. Application to ERP study

For validation 3 (ERP analysis) we contrasted peak amplitudes (calculated on participant-level data) for each of the four methods of cleaning data (e.g., iMARA, manual cleaning, MARA, ‘raw’) (see Figure 3). In the SM section 1.7 we present the same analysis, but with time-frequency analyses rather than ERPs.

**Fig. 3.**
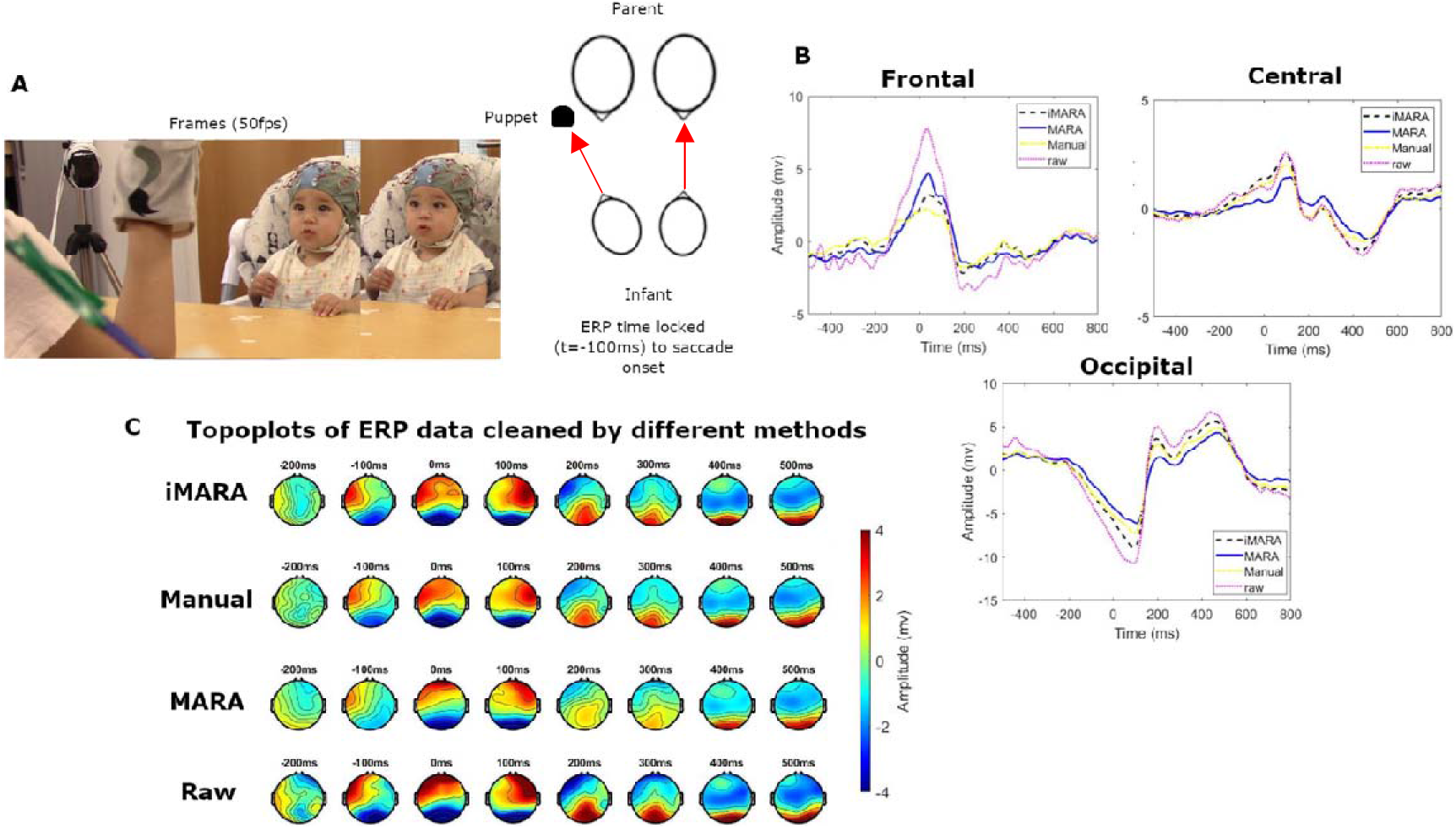
Application of different ICA classification systems to ocular artifact correction in a visual processing ERP study. A) Two-sample frames from which the time-locked gaze shift (−100ms) were identified, and a schematic showing the experimental set up in which mothers were asked to perform a puppet show with their infants. B)Grand average ERPs over frontal pole, central and occipital scalp regions. Different lines show data cleaned by the different systems, e.g., iMARA-retrained infant classifier, MARA-original classifier, Manual classification and also uncleaned ‘raw’ data. C)Topoplots of ERP amplitudes, comparing the different cleaning methods to the raw data.

We used the Tukey procedure to correct for multiple comparisons in the ERP comparisons. Summary tables for all ANOVAs can be found in SM1.1 Table 1. Results from the one-way ANOVAs revealed that peak amplitudes for frontal pole ERPs in the −100 to 100ms time window were significantly lower for all ICA cleaning methods as compared to the raw un-ICA cleaned data. Peak amplitudes for iMARA were lower than for MARA, indicating that more of the ocular artifact had been removed, but this difference was not significant after correcting for multiple comparisons. For central and occipital ERPs, peak amplitudes for MARA were lower than those observed following manual cleaning and cleaning with iMARA, indicating that MARA had removed more genuine neural data. This effect was significant when examining the relationship between MARA and the raw data, but the difference between MARA and iMARA was not significant after correcting for multiple comparisons (p=.10/.11 for central/occipital).

## 4. Discussion

We retrained the popular MARA system for binary (i.e., neural or artifact) classification of ICA-components, to be more sensitive to the types of stereotypical artifacts produced during naturalistic EEG recordings acquired from infants. Our retrained ‘iMARA’ classifier classified artifactual and neural ICA-components from samples of infant EEG with significantly greater levels of agreement with expert manual classification than the original MARA classifier. We examined how well iMARA’s performance generalised to an additional blind dataset as well as its ability to remove ocular-related artifacts in a simple ERP study. Through this, we aimed to provide a tool for developmental EEG researchers wanting to implement automatic ICA cleaning.

### 4.1. Summary of retrained classifier’s performance

In our first validation study, we tested MARA’s and iMARA’s performance against ICA-components manually classified by an expert rater on the full n=44 dataset. Here iMARA achieved a mean classification error rate of 26% (24% with outliers removed), performing significantly better than MARA with a mean error rate of 38%. In the second validation, we tested iMARA on an unseen dataset, collected using the same experimental setup. In this second validation study, iMARA achieved a mean classification error rate of 25%, again significantly outperforming MARA at 38%. Overall, the differences between iMARA and MARA’s agreement with the manual classification and the inter-rater agreement between humans were marginal (7-8% lower average agreement for automatic classification) relative to the overall error rates of either system (25% MSE for automatic and 18% for manual). This is consistent with the error rates between classifier-human and human-human in previous studies (e.g., 5-6% in Winkler et al., 2011). Our retrained iMARA classifier provides, therefore, a more suitable alternative for classifying paediatric ICA-components than the current ‘gold standard’ of manual classification. Additionally, as manual cleaning relies on a large degree of familiarity with ICA and EEG data generally, less experienced researchers using this tool can gain insight into the types of ICA components that are commonly identified as artifacts in paediatric EEG data.

### 4.2. Application of classifiers performance in ERP study

We also compared the performance of the iMARA and MARA to manual classification in a simple ERP study. We examined how well each classifier was able to clean the ERP data, focusing in particular on the removal of activity over frontal pole electrodes at the onset of a saccade (gaze shift) and activity over occipital electrodes after a gaze fixation. Our analysis indicated that all methods of ICA cleaning removed statistically similar amounts of frontal pole activity from the raw (un-ICA-cleaned) data, but that neither the data cleaned manually nor iMARA removed all of the frontal pole activity associated with the eye movement artifact. This is consistent with previous research on adults, which found that standard ICA cleaning methods do not entirely remove all frontal EEG activity associated with eye movement artifacts (Plöchl et al., 2012). This is an important point which should be born in mind in interpreting the results of EEG studies.

Results of validation 3 also show that the post-fixation (gaze onset) visual responses (ERPs) were lower in data cleaned using MARA than for the other types of cleaning, indicating that, while the original MARA classifier did successfully remove comparable amounts of the ocular artifact, it also removed significant amounts of the visually evoked potential (neural signal of interest). This is supported by further analyses (see SM section 1.4, Table S4) which showed that on average MARA removed 64% of components compared to iMARA which removed 39% suggesting that MARA removed more of the total EEG variance. This effect was observed less strongly in the iMARA group, indicating that iMARA had retained more of the original signal than MARA, but this effect was not significant after correcting for multiple comparisons.

### 4.3. Limitations of the current study

There are two explanations for the higher error rates obtained with the current dataset (compared to error rates achieved by Winkler and colleagues 2011). First, the classification of ICA components is notably poorer when applied to lower density electrode montages. In a follow-up study, Winkler and colleagues (2014) found using the original MARA classifier that classification error rates increased from 9% to 32% when comparing 104 to 16 channel electrode setups (although for 32 channel setups it was still comparably lower ~13%) (see also SM section 1.6). This is likely due to the worsening performance of the current density norm feature with lower density setups as this feature relies on estimations of source activity and use of algorithms that are generally only recommended and applied on higher (>64) density electrode setups.

The second reason for the poorer performance compared to previous applications could be due to the increased ambiguity when classifying ICA-components from infant compared to adult EEG. This may be one of the reasons why ICA is not as widely applied within paediatric EEG research as it is within adult EEG research. In our data, we found that averaged across multiple independent coders, infant source components could only be classified with an inter-coder error rate of 18%, compared with 15% for adult data. Similar rates were also achieved when we asked the same coder (coder 1) to classify the same samples of ICA-components at a later time point. Here the agreement between coder 1 (first and second time rating the same 384 infant ICA components) was 17%. Therefore, we suggest that ICA-components from infant EEG (particularly recorded using naturalistic paradigms) are fundamentally more ambiguous because they are more likely to contain a mixture of neural data and artifact, and thus are more difficult to classify binarily.

### 4.4. Recommendations for future research

Future research might explore the iMARA’s ability to separate artifact and neural signals at different frequencies. For example, In SM 1.7 we explore the time-frequency properties of the ERP-responses shown in classifier validation 3. From these plots, it is clear that the classifiers are removing (with varying success) signal that is broadband (i.e., not frequency specific). This may be interesting for future research to explore as eye movements are commonly characterised in time or topographically, but are less often characterised in time-frequency space. Having a clear picture of how ocular artifact in naturalistic data manifests in time-frequency space, as well as, having appropriate tools to identify/ remove it will be of high value to the field going forward. Additionally, it might be useful for future research to integrate iMARA as part of a fully automated EEG pre-processing pipeline either especially for paediatric EEG data or one that is flexible to adult and/or paediatric EEG data.

## 5. Conclusions

This paper presents an automatic ICA classification tool that was specifically tailored to work with infant EEG datasets and EEG data collected during naturalistic parent-infant interactions. We show that the retrained iMARA classifier achieved low classification errors and was better at cleaning stereotypical artifact from a simple visual attention ERP study than the original MARA, adult-trained classifier.

## Acknowledgements

This research was funded by a Project Grant from the Leverhulme Trust, number RPG-2018-281. We wish to thank Federica Lamagna and Martina Eliano for contributing to coding the data. We also wish to thank all families who participated in the research.

## Competing Interests

The authors declare no competing interests.

## Implementation

All MATLAB functions to run the retrained classification system can be found here [https://github.com/Ira-marriott/iMARA/tree/main]

## Notes

https://github.com/Ira-marriott/iMARA/tree/main

### Competing Interest Statement

The authors have declared no competing interest.

## References

Bigdely-Shamlo, N., Mullen, T., Kothe, C., Su, K. M., & Robbins, K. A. (2015). The PREP pipeline: standardized pre-processing for large-scale EEG analysis. Frontiers in neuroinformatics, 9, 16.

Chaumon, M., Bishop, D. V., & Busch, N. A. (2015). A practical guide to the selection of independent components of the electroencephalogram for artifact correction. Journal of neuroscience methods, 250, 47–63.

Cohen, M. X. (2014). Analyzing neural time series data: theory and practice. MIT press.

Cuevas, K., Cannon, E. N., Yoo, K., & Fox, N. A. (2014). The infant EEG mu rhythm: methodological considerations and best practices. Developmental Review, 34(1), 26–43.

Debnath, R., Buzzell, G. A., Morales, S., Bowers, M. E., Leach, S. C., & Fox, N. A. (2020). The Maryland analysis of developmental EEG (MADE) pipeline. Psychophysiology, 57(6), e13580.

Delorme, A., & Makeig, S. (2004). EEGLAB: an open-source toolbox for analysis of single-trial EEG dynamics including independent component analysis. Journal of neuroscience methods, 134(1), 9–21.

Desjardins, J. A., van Noordt, S., Huberty, S., Segalowitz, S. J., & Elsabbagh, M. (2020). EEG Integrated Platform Lossless (EEG-IP-L) pre-processing pipeline for objective signal quality assessment incorporating data annotation and blind source separation. Journal of Neuroscience Methods, 347, 108961.

Dimigen, O. (2020). Optimizing the ICA-based removal of ocular EEG artifacts from free viewing experiments. Neuroimage, 207, 116117.

Farroni, T., Csibra, G., Simion, F., & Johnson, M. H. (2002). Eye contact detection in humans from birth. Proceedings of the National academy of sciences, 99(14), 9602–9605.

Gabard-Durnam, L. J., Mendez Leal, A. S., Wilkinson, C. L., & Levin, A. R. (2018). The Harvard Automated Processing Pipeline for Electroencephalography (HAPPE): standardized processing software for developmental and high-artifact data. Frontiers in neuroscience, 12, 97.

Georgieva, S., Lester, S., Noreika, V., Yilmaz, M. N., Wass, S., & Leong, V. (2020). Toward the Understanding of Topographical and Spectral Signatures of Infant Movement Artifacts in Naturalistic EEG. Frontiers in neuroscience, 14, 352.

Halder, S., Bensch, M., Mellinger, J., Bogdan, M., Kübler, A., Birbaumer, N., & Rosenstiel, W. (2007). Online artifact removal for brain-computer interfaces using support vector machines and blind source separation. Computational intelligence and neuroscience, 2007.

Holleman, G. A., Hooge, I. T., Kemner, C., & Hessels, R. S. (2020). The ‘Real-World Approach’and Its Problems: A Critique of the Term Ecological Validity. Frontiers in Psychology, 11, 721.

Hoormann, J., Falkenstein, M., Schwarzenau, P., & Hohnsbein, J. (1998). Methods for the quantification and statistical testing of ERP differences across conditions. Behavior Research Methods, Instruments, & Computers, 30(1), 103–109.

Jones, E. J., & Johnson, M. H. (2017). Early neurocognitive markers of developmental psychopathology. The Wiley Handbook of Developmental Psychopathology, 197–214.

Jones, E. J. H., Goodwin, A., Orekhova, E., Charman, T., Dawson, G., Webb, S. J., & Johnson, M. H. (2020). Infant EEG theta modulation predicts childhood intelligence. Scientific reports, 10(1), 1–10.

Leach, S. C., Morales, S., Bowers, M. E., Buzzell, G. A., Debnath, R., Beall, D., & Fox, N. A. (2020). Adjusting ADJUST: Optimizing the ADJUST algorithm for paediatric data using geodesic nets. Psychophysiology, e13566.

Lepage, Jean◻François, and Hugo Théoret. “EEG evidence for the presence of an action observation–execution matching system in children.” European Journal of Neuroscience 23.9 (2006): 2505–2510.

Makeig, S., Bell, A. J., Jung, T. P., & Sejnowski, T. J. (1996). Independent component analysis of electroencephalographic data. In Advances in neural information processing systems (pp. 145–151).

Marshall, P. J., Bar-Haim, Y., & Fox, N. A. (2002). Development of the EEG from 5 months to 4 years of age. Clinical Neurophysiology, 113(8), 1199–1208.

McHugh, M. L. (2012). Interrater reliability: the kappa statistic. Biochemia Medica: Biochemia Medica, 22(3), 276–282.

Mullen, T. (2012). CleanLine EEGLAB plugin. San Diego, CA: Neuroimaging Informatics Tools and Resources Clearinghouse (NITRC).

Plöchl, M., Ossandón, J. P., & König, P. (2012). Combining EEG and eye tracking: identification, characterization, and correction of eye movement artifacts in electroencephalographic data. Frontiers in human neuroscience, 6, 278.

Orekhova, E. V., Elsabbagh, M., Jones, E. J., Dawson, G., Charman, T., & Johnson, M. H. (2014). EEG hyper-connectivity in high-risk infants is associated with later autism. Journal of neurodevelopmental disorders, 6(1), 1–11.

Orekhova, E. V., Stroganova, T. A., Posikera, I. N., & Elam, M. (2006). EEG theta rhythm in infants and preschool children. Clinical Neurophysiology, 117(5), 1047–1062.

Piazza, C., Cantiani, C., Miyakoshi, M., Riva, V., Molteni, M., Reni, G., & Makeig, S. (2020). EEG effective source projections are more bilaterally symmetric in infants than in adults. Frontiers in Human Neuroscience, 14.

Risko, E.F., Richardson, D.C., Kingstone, A., 2016. Breaking the Fourth Wall of Cognitive Science: Real-World Social Attention and the Dual Function of Gaze. Curr. Dir. Psychol. Sci. 25, 70–74. https://doi.org/10.1177/0963721415617806

Rutledge, D. N., & Bouveresse, D. J. R. (2013). Independent components analysis with the JADE algorithm. TrAC Trends in Analytical Chemistry, 50, 22–32.

Stroganova, T. A., Orekhova, E. V., & Posikera, I. N. (1999). EEG alpha rhythm in infants. Clinical Neurophysiology, 110(6), 997–1012.

Wass, S. V., Noreika, V., Georgieva, S., Clackson, K., Brightman, L., Nutbrown, R., … & Leong, V. (2018). Parental neural responsivity to infants’ visual attention: how mature brains influence immature brains during social interaction. PLoS biology, 16(12), e2006328.

Wass, S. V., Whitehorn, M., Haresign, I. M., Phillips, E., & Leong, V. (2020). Interpersonal neural entrainment during early social interaction. Trends in cognitive sciences, 24(4), 329–342.

Webb, S. J., Bernier, R., Henderson, H. A., Johnson, M. H., Jones, E. J., Lerner, M. D., … & Westerfield, M. (2015). Guidelines and best practices for electrophysiological data collection, analysis and reporting in autism. Journal of autism and developmental disorders, 45(2), 425–443.

Widmann, A., & Schröger, E. (2012). Filter effects and filter artifacts in the analysis of electrophysiological data. Frontiers in psychology, 3, 233.

Winkler, I., Brandl, S., Horn, F., Waldburger, E., Allefeld, C., & Tangermann, M. (2014). Robust artifactual independent component classification for BCI practitioners. Journal of neural engineering, 11(3), 035013.

Winkler, I., Debener, S., Müller, K. R., & Tangermann, M. (2015, August). On the influence of high-pass filtering on ICA-based artifact reduction in EEG-ERP. In 2015 37th Annual International Conference of the IEEE Engineering in Medicine and Biology Society (EMBC) (pp. 4101–4105). IEEE.

Winkler, I., Haufe, S., & Tangermann, M. (2011). Automatic classification of artifactual ICA-components for artifact removal in EEG signals. Behavioural and brain functions, 7(1), 30.

Xie, W., Mallin, B. M., & Richards, J. E. (2018). Development of infant sustained attention and its relation to EEG oscillations: an EEG and cortical source analysis study. Developmental Science, 21(3), e12562.

Xie, W., Mallin, B. M., & Richards, J. E. (2019). Development of brain functional connectivity and its relation to infant sustained attention in the first year of life. Developmental Science, 22(1), e12703.

