## Supplementary materials for "Automatic classification of ICA components from infant EEG using MARA"

**Supporting Materials**

**Appendix A**

*ICA rejection criteria*

The criteria used for manual ICA classification for infant EEG data were highly similar to the principles suggested for adult EEG data (e.g., Chaumon et al., 2015). Components were marked as artifact/ rejected only under the null hypothesis – when the component is not considered to contain notable amounts of the neural signal. Where a researcher was in doubt over whether a component contains real EEG (neural) we opted to retain that component.

All selection of components was performed using the interface provided by EEGLAB’s *pop_selectcomps* function. Often, EEG researchers only reviewed the first ~10 components, as later components account for little overall variance. The machine-based classifier, however, reviews all components on an individual basis, and so for appropriate comparison coders were asked to rate all components on an individual basis.

Components were judged first on their topography, second on their power spectrum, and third on their time course, according to the flow chart below:

*Flow diagram of human ICA classification:*

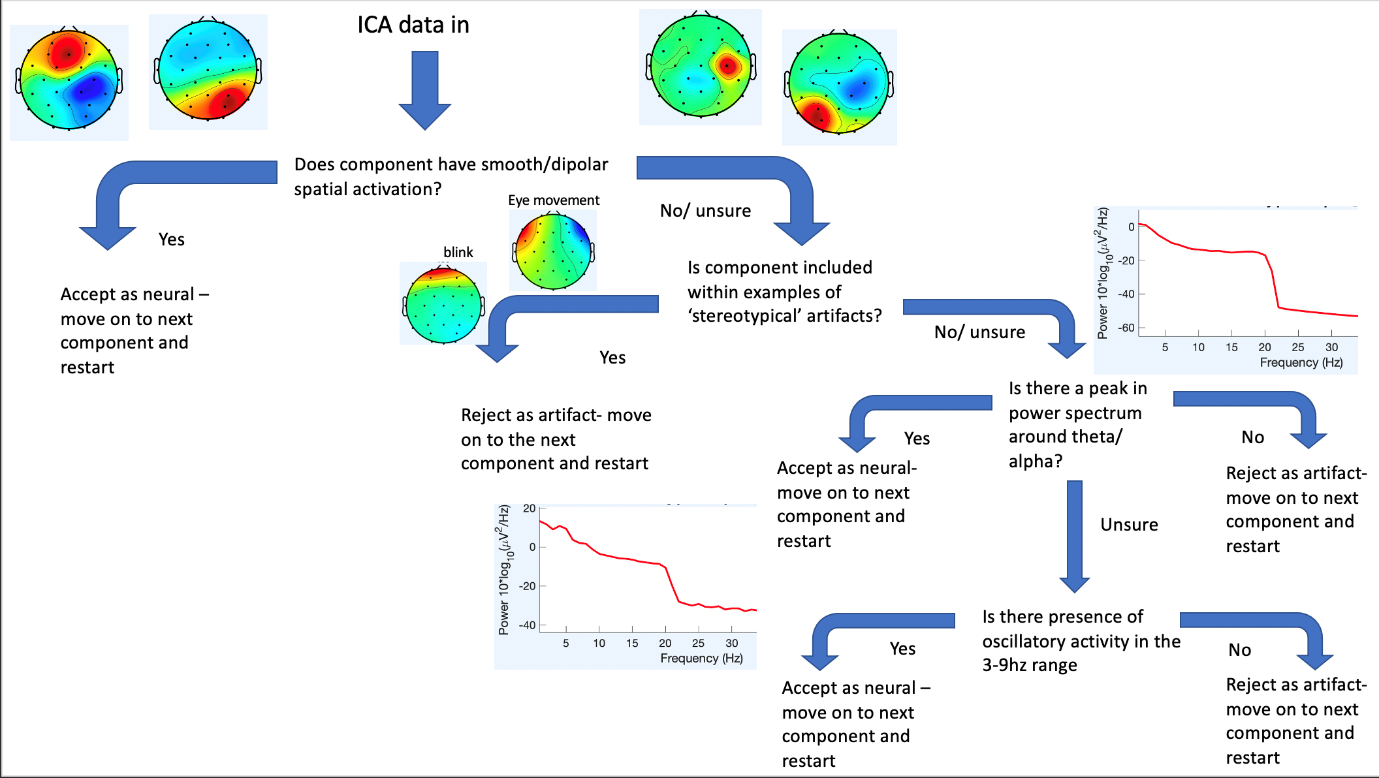

*Stage 1. Evaluation of topographical activation*

Neural components were largely identified by the presence of smooth, dipolar spatial activity. These components were often immediately identifiable in the *pop_selectcomps* component overview and did not need further investigation. A variety of different criteria based on topographical activation were applied:

*Localisation to one electrode.* Components with very localised topographical (i.e., localised to one electrode) activations were always a cause for further investigation (in these cases, coders would move to stage 2). For example:

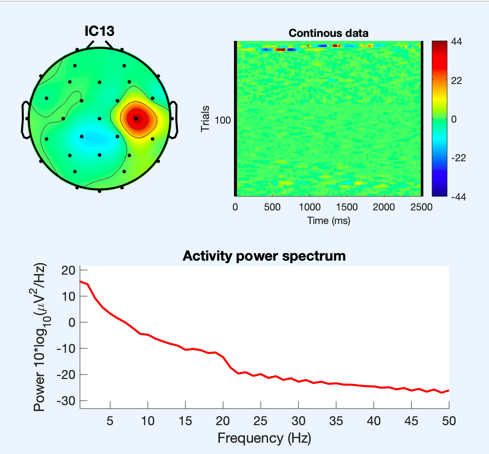

*Frontal peripheral activation.* Certain stereotypical artifactual components were readily identifiable from their topographical maps. For example, components with strong peripheral activations and in particular components with strong/ very localised frontal pole activation (blinks). For example:

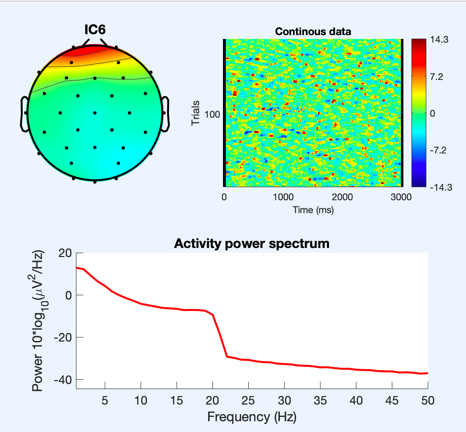

*Bilateral frontal topography.* Components with opposite bilateral frontal topography often indicated horizontal eye movements. For example:

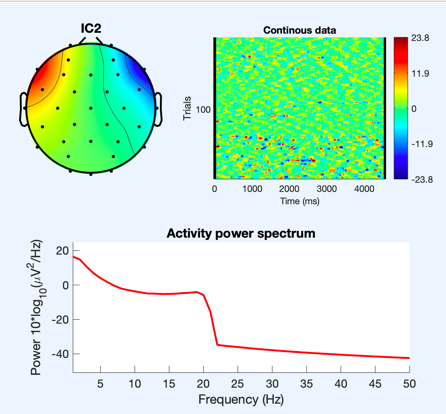

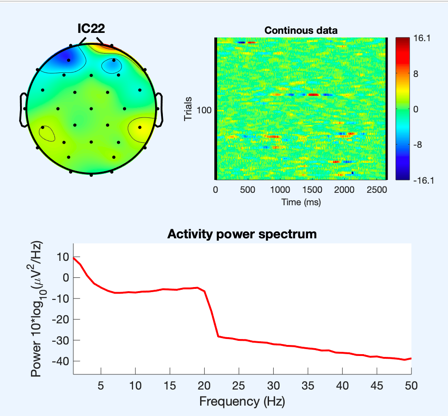

*Temporal peripheral activation*. Components with strong/ very localised temporal activation often indicated jaw/ speech related artefact. For example:

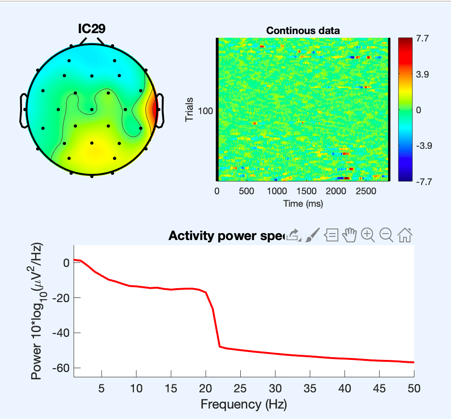

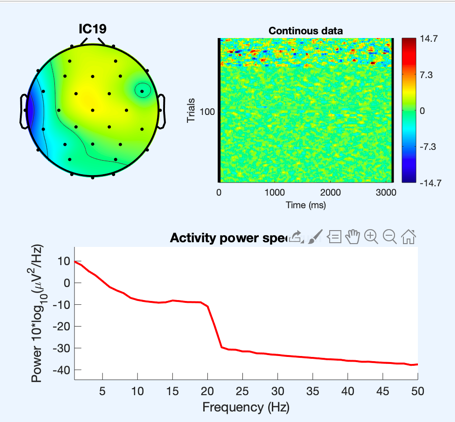

*Activation around P7/P8*. Components with bilateral/strong activation around P7/ P8 (on 10-20 32 channels layout) often indicated neck movement. For example:

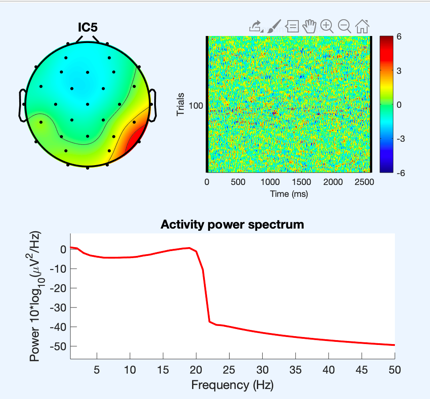

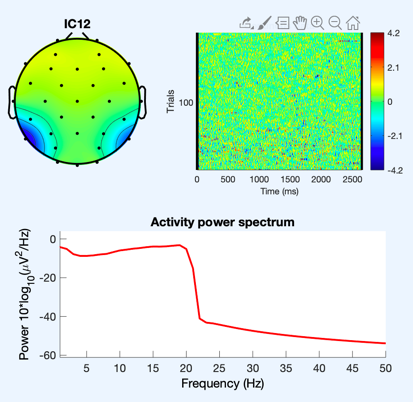

*Stage 2. Evaluation of the power spectrum*

Neural components were identified by 'peaks' in the power spectrum in theta (3-6Hz) and/ or alpha (6-9Hz) range. For example:

 
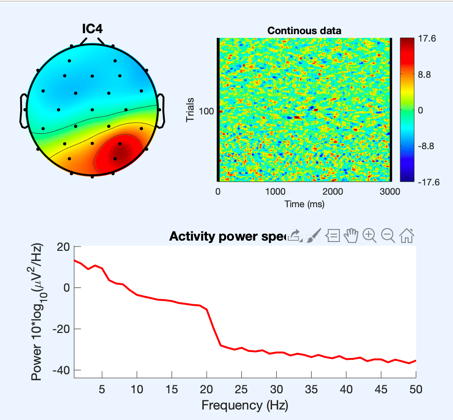

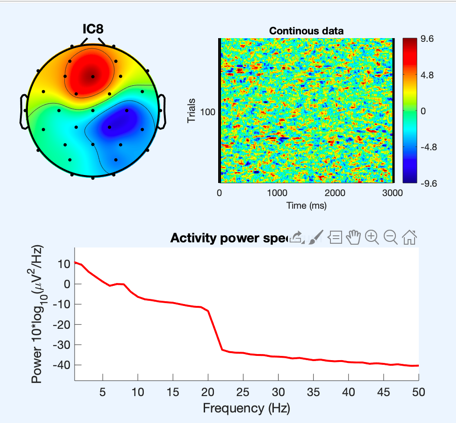

Components with more mixed neural and artefact sources with more subtle peaks in theta/alpha were also identified at this stage as neural. For example:

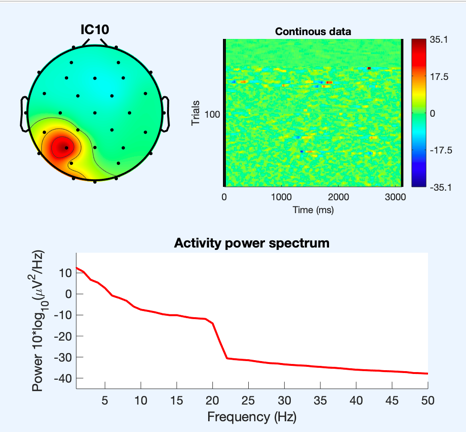

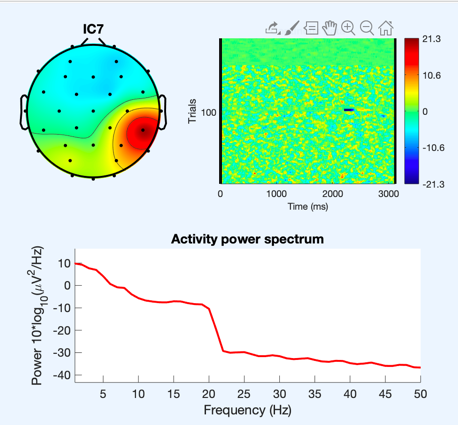

Artefactual components were predominantly identified at this stage through lack of alpha/ theta peak. Further, if a component had no clear 'peak' in alpha but accounted for a lot of total variance coders would move to stage 3 (see below).

Components with power at high frequencies (muscle/ speech artifact) were also marked as artifact at this stage (for examples see the earlier examples for speech and movement-related artifacts).

Components with activity beyond 20Hz were also excluded as the current dataset is low pass filtered at 20Hz – see example below under infrequent high amplitude noise spikes.

*Stage 3. Evaluation of time course*

Artifactual time courses were identified predominantly from their ERP image top right of below figure. If time course activation was mainly driven by infrequent high amplitude spikes (spots of extreme dark colour in ERP image) then it was marked as artefact. For example:

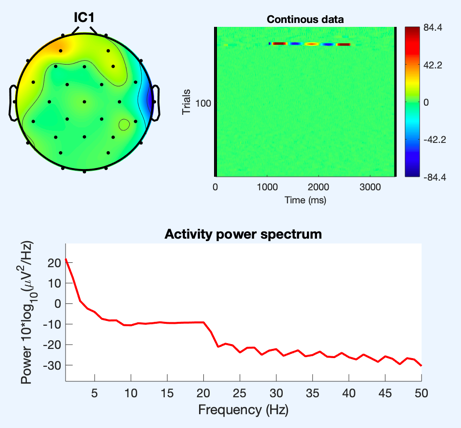

A segment of the component time course:

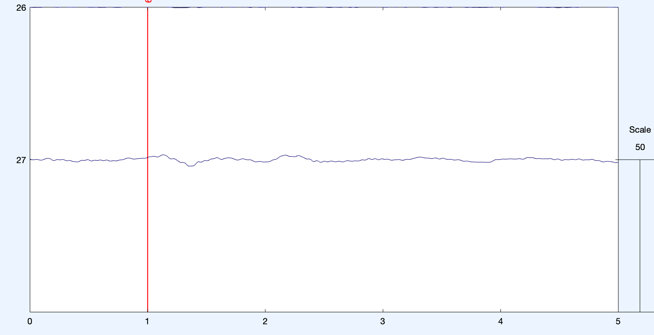

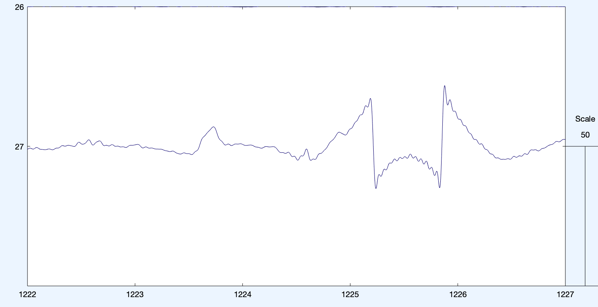

This is different from the example component below as this components time course is predominantly flat throughout.

Other components showed ‘high’ theta/alpha power but no peak. For example:

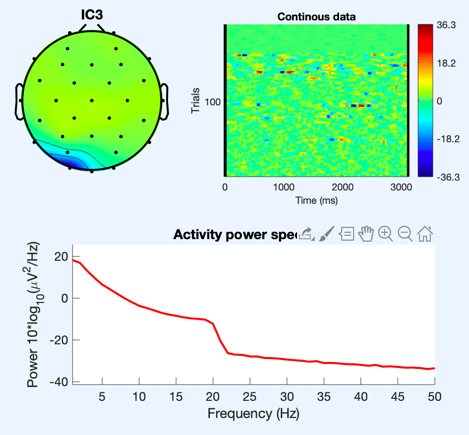

A segment of the component time course:

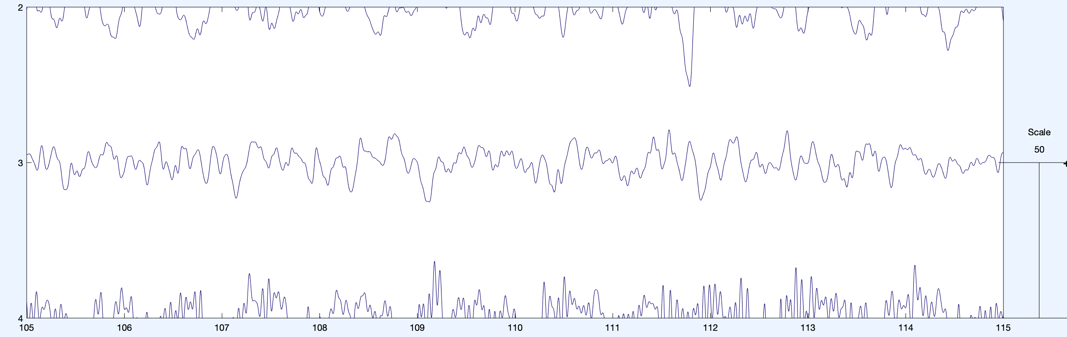

**1. Supplementary Materials**

*1.1 Validation 3: Results of ANOVAs*

*Table S1 summary results of One-Way ANOVA for each scalp region. For frontal pole electrodes, peak amplitudes were compared in the -100 to 100ms time window. For central and occipital components, amplitudes were compared in the 200 to 300ms time window. Electrode groupings used are shown in Table S3.*

| Group ID | | ‘F’ | | | ‘*p’* | | |
| --- | --- | --- | --- | --- | --- | --- | --- |
|  |  | Frontal | Central | Occipital | Frontal | Central | Occipital |
| iMARA | MARA | 2.17 | 2.06 | 6.43**43** | 0.89 | 0.10 | 0.11 |
| iMARA | Manual | 3.92 | 1.54 | 4.39 | 0.85 | 0.70 | 0.88 |
| iMARA | Raw | -2.39 | 0.70 | 1.88 | <0.01 | 0.79 | 0.65 |
| MARA | Manual | 4.74 | 0.57 | 1.39 | 0.42 | 0.60 | 0.41 |
| MARA | Raw | -1.57 | -0.26 | -1.12 | <0.01 | <0.01 | <0.01 |
| Manual | Raw | -3.32 | 0.26 | 0.91 | <0.01 | 0.20 | 0.22 |

*1.2. Inter expert reliability*

*Table S2. Error rates between expert coders on an n=15 subsample of the total data. Each cell shows agreement for infant data (left) and adult data (right)*

|  | Rater 1 | Rater 2 | Rater 3 |
| --- | --- | --- | --- |
| Rater 1 |  | 0.22/0.12 | 0.14/0.15 |
| Rater 2 | 0.22/0.12 |  | 0.19/0.18 |
| Rater 3 | 0.14/0.15 | 0.19/0.18 |  |

*1.3. Electrode positions*

*Table S3. Channel clusters and corresponding 10–20, 32-channel Biosemi positions*

| Clusters | 10-20 Positions | Biosemi 32 channel electrodes |
| --- | --- | --- |
| Frontal pole | Fp1, AF3, AF4, FP2 | 1, 2, 29, 30 |
| Frontal | F7, F3, F4, F8, Fz | 3, 4, 27, 28, 31 |
| Central | C3, CP1, CP5, CP6, CP2, C4, Cz | 5, 6, 8, 9, 10, 21, 22, 23, 25, 26, 32 |
| Temporal | T7, T8 | 7, 24 |
| Parietal | P7, P3, Pz, P4, P8 | 11, 12, 13, 19, 20 |
| Occipital | PO3, O1, Oz, O2, PO4 | 14, 15, 16, 17, 18 |

*1.4. ICA components removed by each method*

*Table S4. Average number and percentage of ICA components removed by each method*

| System | Mean number of components removed (% of total) |
| --- | --- |
| Original MARA | 18 (64%) |
| Retrained MARA | 11 (39%) |
| Hand cleaned | 13 (46%) |

*1.5. 40Hz subsample replication*

All of the data used in this study were narrow-band filtered between 1 and 20Hz. This was done before ICA correction to improve the signal to noise ratios and the quality of the ICA decomposition. However, in their tutorial ([https://irenne.github.io/artifacts/­](https://irenne.github.io/artifacts/%1f)) Winkler and colleagues (2011) note that application of MARA to narrow-band filtered data might result in suboptimal performance, as the spectral features are calculated on the power spectrum between 2 and 39Hz. We, therefore, wanted to test the original classifier’s performance on naturalistic infant data that had not been narrow-band filtered. We processed a subsample of 15 datasets in the same way as described in the main text except for the data was low pass filtered at 40Hz this time instead of 20Hz. We then ran ICA, and all source components were again hand labelled by an expert. Results indicated an MSE between the classifier and human labelling of 23% indicating a good degree of similarity. However, the classifier was removing on average 97% of the approx. 32 components, i.e., retaining on average only one source component. The human labelling also removed 90% of the source components, more than double the amount as the data used in the main text. Whilst the MSE is slightly lower than the retrained classifier in the main text, it is clear that this method is far from optimal as clearly if any method is removing over 95% of the total variance it is also removing large amounts of genuine neural activity.

*1.6. MARA adapted strategy*

In their follow up study in which Winkler and colleagues (2014) tested the robustness of their classifier on a variety of novel experimental paradigms and electrode setups, they found that when applied to data with lower density electrode setups e.g. 16 or 32 channels, the classifier’s error rate increased linearly from an MSE between automatic and manual classification from 9 to 32%. This suggests that the classifier performs significantly worse with lower density electrode setups. The reason for this decrease in performance was due to the spatial features performing notably worse (Winkler et al., 2014). The MSE of the CDN feature on 32 and 16 channel setups rose to over 50% compared to 12% with higher density setups. To improve the performance of the original MARA classifier with lower density EEG recordings, the authors proposed an adaptive strategy in which the classifier is subtly adapted to fit the study-specific electrode montage i.e., re-training the classifier on the patterns cut to the specific electrode setup. In their study application of the adaptive strategy with 16 and 32 channel setups lead to an MSE of approx. 16%. We also tested this adaptive strategy on our 32 channel infant EEG data to see whether this led to a similar error rate. We did this by adapting the electrode montage used for spatial feature identification to the Biosemi 32-channel layout but using the same training data as used by the original MARA classifier. This led to an MSE between the adapted strategy classifier and the human labelling of 43%.

Neither the original classifier nor the adapted strategy performed well when applied to our infant EEG data. On one hand, we have a system that achieved decent rates of agreement with hand labelling i.e., MSE of 23% but removes over 95% of the data (e.g., original MARA); on the other hand, the adaptive strategy removes fewer components but has a much higher error rate. Therefore, the final option is to treat the infant data as distinct, retraining the classifier using infant source components and the most salient features for infant EEG.

*1.7. Time-Frequency analysis of ERP data presented in validation study 3*

To further investigate the removal of ocular artifact reported in validation 3. We examined time-frequency responses for the different methods to assess the time-frequency representations of the signal and how this was affected by the ICA cleaning methods. For this analysis single-trial data were first decomposed into their time-frequency representation by multiplying the power spectrum of the EEG (obtained from the fast-Fourier-transform) by the power spectrum of complex Morlet wavelets [*ei*2π*tfe*−*t*2/(2σ2), where *t* is time, *f* is the frequency (which increased from 2 to 16 Hz in 15 linearly spaced steps), and σ defines the width of each frequency band, set according to σ/(2π*f*), σ was set to scale with increases in the centre frequency of the wavelet. We set this parameter to increase logarithmically from 3-10 cycles in 15 increments], and then taking the inverse fast-Fourier-transform. From the resulting complex signal, an estimate of frequency band-specific power at each time point was defined as the squared magnitude of the result of the convolution *Z* (real[*z*(*t*)]2 + imag[*z*(*t*)]^2^). Power was then decibel normalised to data in the -1000 to -700ms time window.

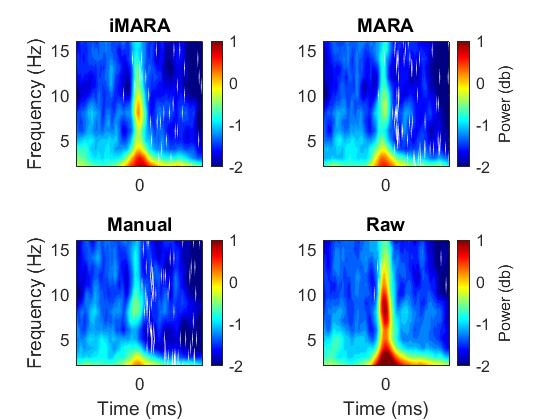

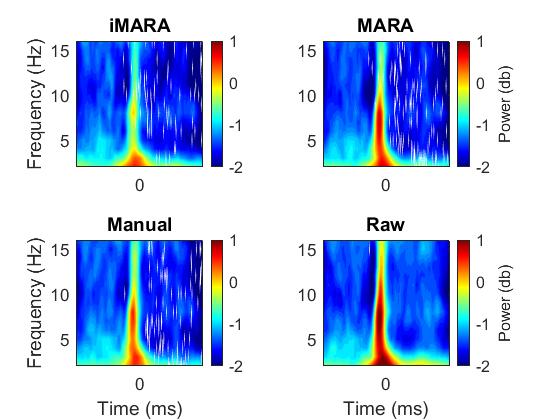

**Frontal Pole TF-Power**

**Occipital TF-Power**

B

A

-2.5	0	2.5

-2.5	0	2.5

-2.5	0	2.5

-2.5	0	2.5

Time (s) (s)

Time (s) (s)

-2.5	0	2.5

Time (s)

-2.5	0	2.5

-2.5	0	2.5

Time (s)

-2.5	0	2.5

Time (s)

Time (s) (s)

Time (s) (s)

*Fig.4 Time-frequency power relative to the onset of infant looks to partner in data cleaned using original 'MARA', retrained 'iMARA’ and manual ICA classification and compared to raw data. A) shows time-frequency plots for data cleaned using 4 methods over frontal pole electrodes. B) shows time-frequency plots for data cleaned using 4 methods over occipital electrodes*

From **Fig. 4** we can see that all methods of cleaning resulted in broadband removal of the signal both frontally and occipitally. We can also see that the different methods are to a varying degree removing most but not all the ocular artifact time-locked to the shift in attention. This may be an interesting possibility for future research to explore as commonly eye movements are characterised in time or space but are less often characterised in time-frequency space.

**References for supplementary materials**

Chaumon, M., Bishop, D. V., & Busch, N. A. (2015). A practical guide to the selection of independent components of the electroencephalogram for artifact correction. *Journal of neuroscience methods*, *250*, 47-63.

Winkler, I., Brandl, S., Horn, F., Waldburger, E., Allefeld, C., & Tangermann, M. (2014). Robust artifactual independent component classification for BCI practitioners. *Journal of neural engineering*, *11*(3), 035013.

Winkler, I., Haufe, S., & Tangermann, M. (2011). Automatic classification of artifactual ICA-components for artifact removal in EEG signals. *Behavioural and brain functions*, *7*(1), 30.
